# Identification and quantification of meat product ingredients by whole-genome metagenomics (All-Food-Seq)

**DOI:** 10.1101/763458

**Authors:** Sören Lukas Hellmann, Fabian Ripp, Sven-Ernö Bikar, Bertil Schmidt, René Köppel, Thomas Hankeln

**Affiliations:** Institute of Organismic and Molecular Evolution, Molecular Genetics & Genome Analysis, Johannes Gutenberg University Mainz, Mainz, Germany; MVZ Labor Volkmann, Karlsruhe, Germany; StarSEQ GmbH, Mainz, Germany; Institute of Informatics, Johannes Gutenberg University Mainz, Mainz, Germany; Official Food Control Authority of the Canton Zurich, Zurich, Switzerland

**Keywords:** Food Metagenomics, Species Identification, Doner Kebap, Read Mapping, Next-Generation-Sequencing

## Abstract

Complex food matrices bear the risk of intentional or accidental admixture of non-declared species. Moreover, declared components can be present in false proportions, since expensive taxa might be exchanged for cheaper ones. We have previously reported that PCR-free metagenomic sequencing of total DNA extracted from sausage samples combined with bioinformatic analysis (termed All-Food-Seq, AFS), can be a valuable screening tool to identify the taxon composition of food ingredients. Here we illustrate this principle by analysing regional Doner kebap samples, which revealed unexpected and unlabelled poultry and plant components in three of five cases. In addition, we systematically apply AFS to a broad set of reference meat material of known composition (i.e. reference sausages) to evaluate quantification accuracy and potential limitations. We include a detailed analysis of the effect of different food matrices and the possibility of false-positive sequence read assignment to closely related species, and we compare AFS quantification results to quantitative real-time PCR (qPCR) and droplet digital PCR (ddPCR). AFS emerges as a potent PCR-free screening tool, which can detect multiple target species of different kingdoms of life within a single assay. Mathematical calibration accounting for pronounced matrix effects can significantly improves AFS quantification accuracy. In comparison, AFS performs better than classical qPCR, and is on par with ddPCR.

## Introduction

The determination and quantification of food ingredients is an important issue in official food control. The complexity of foodstuff, difficulties in the traceability of trading channels and the globalisation of food markets opens doors for fraud and failures in correct labelling, stocking and processing procedures [1]. Possible consequences for consumers are manifold beginning with compliance of ethical aspects like halal, kosher or vegan over health risks caused by pathogenic organisms to simple deception because of economic reasons. In fact, biological contaminants made up the vast majority of warning notices released by the German authorities [2] between 2011 and 2015. The majority of these cases were provoked by microbiological contaminations or the presence of non-declared allergenic food components. Therefore, food and drug legislation demands proper declaration of ingredients and compliance to storage and transport conditions [3, 4]. To ensure adherence to law and to maintain consumer’s safety, there is a growing need for methods that allow for precise determination of food ingredients, ideally spanning all kingdoms of life including plants, animals, bacteria, fungi and perhaps also extending to viruses. A broad palette of analytical methods for analysing foodstuffs has been developed and is routinely applied at official food control laboratories, but also private and industrial control labs. Among these, DNA-based methods like PCR are probably the most widely used technologies, because of their high sensitivity and the possibility to perform quantitative measurements [5–13]. However, even when multiplexed or performed in the meta-barcoding format, PCR-based approaches have the drawback to detect only a limited range of target species and produce assay-dependent amplification biases [14–17].

We have previously shown that deep metagenomic DNA sequencing of whole-genome DNA from foodstuffs, followed by dedicated bioinformatic analysis, is in principle able to overcome these issues. DNA sequence reads obtained from food can be bioinformatically assigned to existing reference genomes for species identification, and the number of reads successfully assigned to a respective genome can be counted to give a quantitative measure of the species proportions. Importantly, such whole-genome sequencing of foodstuff DNA (termed All-Food-Seq: AFS; [18, 19]) does not require any *a priori* definition of possible target species. AFS can therefore be viewed as a screening method, which theoretically can detect an infinite spectrum of diverse species, being only limited by our current knowledge of genomes, as represented in the fast-growing public sequence databases. The “identification plus quantification” principle based on read-assignment and read-counting has been successfully demonstrated so far as a proof-of-principle in a limited number of foodstuff samples, i.e. sausages of pre-defined composition prepared as reference material [8, 20]. We therefore decided to further investigate the potential of AFS in a real-life test case, analysing different doner kebap samples obtained from snack bars. We also saw the necessity to evaluate the quantification potential of AFS in more detail. Inferring species proportions from DNA read proportions can be difficult because it may substantially depend on the food composition and processing. As an example, high quality meat may be substituted for in a product by the addition of rind, lard or skin, which could affect DNA amounts per gram tissue, and thus the inference of species proportions within the foodstuff. To study this so-called matrix effect, we have applied the AFS method to an extended set of reference sausage samples, each containing known admixtures of different meat sources, but prepared according to different recipes [8, 20]. We compared the AFS quantification results to those obtained by qPCR and ddPCR on the same samples and evaluated the effects of matrix composition.

## Materials and Methods

### Food samples and DNA extraction

Doner kebap samples were purchased at five snack bars distributed in the Rhine-Main area. All meat pieces were identified by eye and selected by sterile forceps for subsequent homogenization in large volume using a standard kitchen device. About 1 g of the homogenized matrix, which looked surprisingly different (ranging between an oily and granular texture), was taken for subsequent DNA isolation using the Wizard Plus Miniprep DNA purification system (Promega, Madison, USA) according to the manufacturer’s protocol. DNA was quantified by Qubit fluorometry (Thermofisher Scientific, Schwerte, Germany).

Calibration sausage samples containing admixtures of cattle, chicken, pig, sheep and turkey at defined amounts were produced by a professional butchery and provided by the Official Food Control Authority of the Canton Zürich, Switzerland [8, 20]. The samples were prepared for calibration of foodstuff detection methods and reflect three different recipes of sausage production (Online Tab. S1): AllMeat sausage (Kal A-E: meat), Lyoner-style sausage (KLyo A-D: matrix of meat, rind and lard) and Poultry-Lyoner (KGeflLyo A-D: matrix of meat and skin). Total DNA was extracted out of 200 mg homogenized sausage sample using the Wizard Plus system (Promega, Madison, USA) according to the manufacturer’s protocol.

### Illumina library preparation and sequencing

Sequencing library preparations and sequencing were performed by a commercial provider (StarSEQ, Mainz, Germany). The Nextera DNA Library Preparation Kit (Illumina, San Diego, USA) was applied following the manufacturer’s instructions. Typically, 1 ng of total DNA was used. Sequencing was carried out on an Illumina MiSeq instrument using reagent kit v.2 in 150 bp paired-end (reference sausages) and 50 bp single-end (doner kebap samples) mode, respectively. In principle, both sequencing modes deliver comparably valid results [19]. Between 200 k and 2600 k reads were generated per sample (Online Tab. S1). Adjustments by downsampling were omitted, because our previous analysis showed that read numbers > 100 k produced consistent quantification results independent of dataset size [19]. All datasets were quality checked, trimmed and filtered by using FASTQC data evaluation software [21] and trimmomatic v0.33 trimming tool [22]. Datasets have been submitted to the SRA database under the project names PRJNA271645 and PRJEB34001.

### Bioinformatic analysis of main ingredients using AFS

The AFS read-mapping pipeline was executed with 3 rounds of iterative mapping and step-wise decreased mapping stringency, as described [18, 19].This strategy allows for a final number of 2 mismatches after mapping step 3. At each round, reads that mapped against one of the provided reference genomes were cumulatively counted and reported on a 1-100% scale to reflect relative species proportions. In the doner kebap screening analysis, sequence reads were mapped against a selection of reference genomes (accession numbers: cattle: NC_037328.1, sheep: NC_040252.1, goat: NC_030808.1, pork: NC_010443.5, horse: NC_009144.3, chicken: NC_006088.5, turkey: NC_015011.2, maize: NC_024459.2 and soy: NC_016088.3. In the quantification analysis of the calibrator sausages, reference genome choice was limited to the animal species cattle, chicken, pig, horse, sheep, goat, water buffalo (accession number: NC_037545.1) and turkey. Goat and water buffalo genomes were added to test the robustness of AFS towards false positive signals to be expected between closely related species. All evaluations were performed on a standard desktop PC (Intel(R) Core(TM) i7-8700 CPU @ 3.20GHz, 16GB DDR4 2667 MHz RAM, 256GB SATA SSD, CentOS Linux release 7.6.1810).

Reads that did not match, very likely originate from species not provided as a reference during the AFS mapping step. These unmapped reads (around 3 % per sample), often representing spice plants and microbiota [18], did not undergo further metagenomic analysis in the present study, since the prime goal was to evaluate the quantification properties of AFS for the main meat components.

### Calculation of false-positive read assignments

In order to determine false-positive read assignment rates for the tested species, in particular the closely related cattle-buffalo, chicken-turkey and goat-sheep, we created *in silico* datasets of different proportions of reads for each species with the corresponding related species being absent. To this end, we used whole-genome shotgun datasets from the SRA (SRR8588004, SRR9663406, SRR8442931, SRR8560982, SRR6470934) and performed data pre-processing as described for the reference sausages. For each species, proportions of 1, 5, 10, 25, 50, 75 and 100 % were extracted using the reformat tool from the BBMap suite [23] and complemented to 1 mio reads with the non-related plant species rice (accession number: NC_008394.4). For the cattle-buffalo species pair, we only inspected the false-positive rate of buffalo assignments given a cattle ingredient, as the opposite direction is irrelevant to food safety inspections in our opinion. To investigate the effect of sequence read length on false positive mapping, all generated datasets were trimmed using the reformat tool to a length of 50, 100 and 150 bp, respectively. Subsequent AFS analyses were performed as described above with 3 mapping rounds (accepting max. 2 mismatches) against buffalo, cattle, chicken, goat, horse, pork, sheep and turkey genomes.

## Results

### AFS screening of species composition in doner kebap samples

Doner kebap samples were obtained from five snack bars in the Rhine-Main area. Their meat components were sampled, homogenized and the extracted DNA sequenced. AFS analysis revealed that samples 2 and 3 were prepared from pure beef, while samples 1, 4 and 5 consisted of beef and turkey, with the latter as the dominant component (Tab. 1a). Samples 1, 3 and 5 revealed measurable amounts of soybean DNA (0.5-0.8 %), and sample 1 additionally contained maize DNA (1.8 %). In samples 2, 3 and 5, we observed that 0.1-0.4 % of sequence reads were assigned to goat and sheep. Since the latter also belong to the family of Bovidae, one could interpret the goat/sheep read assignments as candidate false-positives, produced as a consequence of the phylogenetic relatedness and the presence of conserved genomic elements. However, our detailed evaluation of possible false-positive values (see below; Online Fig. S1b, Online Tab. S2) shows that at least for samples 2 and 3 the measured values of goat and sheep are slightly higher than expected for a matrix consisting almost only of cattle. We therefore cannot rule out that small amounts of sheep and goat material were indeed present in these doner samples, allegedly caused by the presence of cheese matrices or due to unknown circumstances during doner production. In contrast, the 0.2 and 0.3 % of chicken reads in samples 1 and 4, which are clearly dominated by turkey, may accordingly be considered false-positives.

**Tab. 1.** Quantification results of AFS pipeline. Raw results obtained by AFS analysis as well as calibrated AFS values obtained by linear regression are compared to PCR-based quantification for **(a)** Doner Kebab samples, **(b)** Kal A-E, **(c)** Klyo A-D and **(d)** KGeflLyo A-D. qPCR data were obtained from [8, 20], ddPCR data from [13]. “Sum dev” represents the sum of % deviation from expected proportions. Best results for each sausage are shaded grey. (n.a. = not analysed)

|  | beef | turkey | soy | maize | horse | pork | chicken | sheep | goat |
| --- | --- | --- | --- | --- | --- | --- | --- | --- | --- |
| Doner Kebab 1 | 8.6 | 88.6 | 0.7 | 1.8 | 0.0 | 0.0 | 0.2 | 0.0 | 0.0 |
| Doner Kebab 2 | 99.5 | 0.0 | 0.0 | 0.0 | 0.0 | 0.0 | 0.0 | 0.2 | 0.3 |
| Doner Kebab 3 | 98.7 | 0.0 | 0.4 | 0.0 | 0.0 | 0.0 | 0.0 | 0.4 | 0.4 |
| Doner Kebab 4 | 5.1 | 94.5 | 0.0 | 0.0 | 0.0 | 0.0 | 0.3 | 0.0 | 0.0 |
| Doner Kebab 5 | 27.7 | 71.3 | 0.4 | 0.0 | 0.0 | 0.0 | 0.3 | 0.1 | 0.1 |

**(b)**
|  |  | beef | pork | sheep | horse | chicken | turkey | goat | buffalo | sum dev |
| --- | --- | --- | --- | --- | --- | --- | --- | --- | --- | --- |
| Kal A | expected | 1.0 | 35.0 | 9.0 | 55.0 | 0.0 | 0.0 | 0.0 | 0.0 |  |
|  | AFS | 1.4 | 30.9 | 10.1 | 57.3 | 0.0 | 0.0 | 0.3 | 0.0 | 8.2 |
|  | AFS cal. | 0.3 | 34.9 | 10.3 | 54.3 | 0.0 | 0.0 | 0.3 | 0.0 | 3.1 |
|  | qPCR | 0.4 | 39.3 | 8.9 | 51.5 | n.a. | n.a. | n.a. | n.a. | 8.6 |
| Kal B | expected | 9.0 | 55.0 | 1.0 | 35.0 | 0.0 | 0.0 | 0.0 | 0.0 |  |
|  | AFS | 11.2 | 49.5 | 1.3 | 37.8 | 0.0 | 0.0 | 0.1 | 0.1 | 11.0 |
|  | AFS cal. | 9.6 | 54.2 | 1.0 | 35.1 | 0.0 | 0.0 | 0.1 | 0.1 | 1.7 |
|  | qPCR | 23.0 | 51.0 | 1.5 | 24.4 | n.a. | n.a. | n.a. | n.a. | 29.1 |
| Kal C | expected | 25.0 | 25.0 | 25.0 | 25.0 | 0.0 | 0.0 | 0.0 | 0.0 |  |
|  | AFS | 25.8 | 19.8 | 28.2 | 25.1 | 0.0 | 0.0 | 1.0 | 0.2 | 10.5 |
|  | AFS cal. | 23.7 | 22.4 | 29.0 | 23.7 | 0.0 | 0.0 | 0.9 | 0.2 | 10.3 |
|  | qPCR | 34.0 | 18.8 | 25.8 | 21.4 | n.a. | n.a. | n.a. | n.a. | 19.6 |
|  | ddPCR | 25.6 | 25.3 | 24.6 | 24.5 | n.a. | n.a. | n.a. | n.a. | 1.8 |
| Kal D | expected | 35.0 | 9.0 | 55.0 | 1.0 | 0.0 | 0.0 | 0.0 | 0.0 |  |
|  | AFS | 38.2 | 7.7 | 50.8 | 1.3 | 0.0 | 0.0 | 1.7 | 0.3 | 11.1 |
|  | AFS cal. | 35.3 | 9.1 | 52.1 | 1.5 | 0.0 | 0.0 | 1.7 | 0.3 | 5.8 |
|  | qPCR | 29.9 | 7.6 | 61.7 | 0.9 | n.a. | n.a. | n.a. | n.a. | 13.3 |
| Kal E | expected | 55.0 | 1.0 | 35.0 | 9.0 | 0.0 | 0.0 | 0.0 | 0.0 |  |
|  | AFS | 56.9 | 1.0 | 32.2 | 8.5 | 0.0 | 0.0 | 1.1 | 0.4 | 6.7 |
|  | AFS cal. | 54.5 | 1.9 | 33.8 | 8.4 | 0.0 | 0.0 | 1.1 | 0.4 | 4.7 |
|  | qPCR | 51.2 | 1.0 | 37.2 | 10.5 | n.a. | n.a. | n.a. | n.a. | 7.5 |

(c)
|  |  | beef | pork | sheep | horse | chicken | turkey | goat | buffalo | sum dev |
| --- | --- | --- | --- | --- | --- | --- | --- | --- | --- | --- |
| **KLyo A** | expected | 14.0 | 80.0 | 0.0 | 0.0 | 0.5 | 5.5 | 0.0 | 0.0 |  |
|  | AFS | 19.3 | 74.6 | 0.0 | 0.0 | 0.7 | 5.3 | 0.0 | 0.1 | 11.2 |
|  | AFS cal. | 13.0 | 80.8 | 0.0 | 0.0 | 0.3 | 5.7 | 0.0 | 0.1 | 2.3 |
|  | ddPCR | 14.2 | 80.2 | n.a. | n.a. | 0.4 | 5.3 | n.a. | n.a. | 0.7 |
| **KLyo B** | expected | 36.0 | 58.0 | 0.0 | 0.0 | 2.0 | 4.0 | 0.0 | 0.0 |  |
|  | AFS | 41.8 | 52.4 | 0.0 | 0.0 | 2.2 | 3.4 | 0.0 | 0.3 | 12.5 |
|  | AFS cal. | 35.9 | 57.9 | 0.0 | 0.0 | 2.1 | 3.8 | 0.0 | 0.3 | 0.8 |
|  | ddPCR | 34.9 | 60.1 | n.a. | n.a. | 1.4 | 3.7 | n.a. | n.a. | 4.1 |
| **KLyo C** | expected | 58.0 | 36.0 | 0.0 | 0.0 | 4.0 | 2.0 | 0.0 | 0.0 |  |
|  | AFS | 66.0 | 28.3 | 0.0 | 0.0 | 3.9 | 1.4 | 0.0 | 0.4 | 17.0 |
|  | AFS cal. | 60.6 | 33.1 | 0.0 | 0.0 | 4.1 | 1.7 | 0.0 | 0.4 | 6.3 |
|  | ddPCR | 58.4 | 36.8 | n.a. | n.a. | 2.9 | 1.9 | n.a. | n.a. | 2.4 |
| **KLyo D** | expected | 80.0 | 14.0 | 0.0 | 0.0 | 5.5 | 0.5 | 0.0 | 0.0 |  |
|  | AFS | 82.8 | 11.3 | 0.0 | 0.0 | 5.0 | 0.4 | 0.0 | 0.5 | 6.8 |
|  | AFS cal. | 77.7 | 15.7 | 0.0 | 0.0 | 5.3 | 0.7 | 0.0 | 0.5 | 4.9 |
|  | ddPCR | 79.2 | 15.9 | n.a. | n.a. | 4.3 | 0.6 | n.a. | n.a. | 4.0 |

(d)
|  |  | beef | pork | sheep | horse | chicken | turkey | goat | buffalo | sum dev |
| --- | --- | --- | --- | --- | --- | --- | --- | --- | --- | --- |
| **KGefiLyo A** | expected | 0.5 | 5.5 | 0.0 | 0.0 | 14.0 | 80.0 | 0.0 | 0.0 |  |
|  | AFS | 0.6 | 5.1 | 0.0 | 0.0 | 29.8 | 64.5 | 0.0 | 0.0 | 31.9 |
|  | AFS cal. | 0.2 | 5.6 | 0.0 | 0.0 | 10.0 | 84.1 | 0.0 | 0.0 | 8.5 |
|  | ddPCR | 0.8 | 9.4 | n.a. | n.a. | 24.6 | 65.1 | n.a. | n.a. | 29.7 |
| **KGefiLyo B** | expected | 2.0 | 4.0 | 0.0 | 0.0 | 36.0 | 58.0 | 0.0 | 0.0 |  |
|  | AFS | 2.0 | 3.5 | 0.0 | 0.0 | 56.9 | 37.6 | 0.0 | 0.0 | 41.9 |
|  | AFS cal. | 2.2 | 3.9 | 0.0 | 0.0 | 40.3 | 53.5 | 0.0 | 0.0 | 9.1 |
|  | ddPCR | 2.1 | 5.0 | n.a. | n.a. | 41.7 | 51.2 | n.a. | n.a. | 13.6 |
| **KGefiLyo C** | expected | 4.0 | 2.0 | 0.0 | 0.0 | 58.0 | 36.0 | 0.0 | 0.0 |  |
|  | AFS | 3.4 | 1.4 | 0.0 | 0.0 | 75.7 | 19.5 | 0.0 | 0.0 | 35.4 |
|  | AFS cal. | 4.4 | 1.8 | 0.0 | 0.0 | 61.2 | 32.6 | 0.0 | 0.0 | 7.2 |
|  | ddPCR | 4.2 | 2.4 | n.a. | n.a. | 62.6 | 30.8 | n.a. | n.a. | 10.4 |
| **KGefiLyo D** | expected | 5.5 | 0.5 | 0.0 | 0.0 | 80.0 | 14.0 | 0.0 | 0.0 |  |
|  | AFS | 3.9 | 0.4 | 0.0 | 0.0 | 89.2 | 6.5 | 0.0 | 0.0 | 18.4 |
|  | AFS cal. | 5.2 | 0.7 | 0.0 | 0.0 | 76.4 | 17.8 | 0.0 | 0.0 | 7.9 |

### AFS quantification of meat ingredients in reference sausages

To specifically study the quantification properties of AFS in a broad set of samples, a total of 13 reference sausage samples (Online Tab. S1), prepared according to three different standard recipes, were sequenced and analysed. Datasets were then studied to evaluate quantification accuracy, the impact of different matrices (i.e. meat, rind, lard and skin), and the probability of false-positive read assignments. AFS results were then compared to quantification data previously obtained by qPCR [8, 20] and droplet digital (dd) PCR [13] on the very same sausage samples (Tab. 1b-D).

#### AFS quantification accuracy

Our sample set covered expected species proportions from 0.5-80 %. Minimal and maximal expected components varied between the three different sample types (meat-only samples Kal A-E 1-55%, mixed-matrix samples KLyo A-D and KGeflLyo A-D 0.5-80%). It turned out that even the low concentrations of ingredients could be detected by AFS with high accuracy (0.5 ± 0.1 %; 1 ± 0.1 %; 2 ± 0.4 %; 4 ± 0.2 %; 5.5 ± 0.5 %). As species concentrations increased, absolute deviations of measured values also increased to a maximum of 20.9 % for the 9-36 % interval and 20.4 % for the 55-80 % interval, respectively (Tab. 1b-D). To compare the performance of AFS for the different sausage types, we summed up the individual species deviations for each sausage individually. Results showed that, omitting any calibration calculations (see below), Kal A-E samples were quantified with overall best results (ranging between 6.7 and 11.1 % deviation), followed by KLyo A-D (6.8 to 17.0 %) and KGeflLyo A-D (18.4 to 41.9 %).

#### Evaluation of false-positive read assignments between related species

The species assignment of sequence reads in AFS is based on classical read-mapping algorithms involving sequence alignment [18]. This implies the potential danger of a mis-classification if a read contains highly conserved DNA sequences, often present in the genomes of phylogenetically closely related taxa. Of course such false positive assignments could have, if present, an eminent effect on detection accuracy. To evaluate the potential of such false-positive read assignment within AFS, we intentionally included in the read-mapping step the reference genomes of species, which are <u>not</u> present in the sausages, but which are evolutionarily close to the real food components. Specifically, we added the genome of the water buffalo (*Bubalus bubalis*), which shared a common ancestor with cattle 13 mio years ago, and the goat (*Capra hircus*), which diverged from sheep 10 mio years ago ([24]; Online Fig. S1a). False-positive signals of buffalo and goat from sausages containing cattle or sheep as real ingredients ranged between 0.0-1.7 % and depended on the amount of the corresponding real ingredient species (Tab. 1b-D). For example, the maximal value of 1.7 % false goat reads was obtained for the Kal D sausage containing 55% sheep.

To systematically specify the chance of false-positive read assignments between species pairs in AFS, we simulated read datasets with varying, known amounts of reads from the species in our study and mapped them to the respective reference genomes. The amount of false-positive reads in fact scaled linearly with the real ingredient proportions (Online Fig. S1b, Online Tab S2), allowing us to define threshold values for the respective species pairs. Interestingly, but not unexpectedly, the short 50 bp reads produced markedly higher false-positive values than 100 bp and 150 bp reads. For example, a 100% sheep dataset produced 5.1% false-positive goat assignments with 50 bp read length, but only 2.7 % with 150 bp reads (Online Fig. S1b). Some minor ‘asymmetric’ quantification results (i.e. chicken against turkey genome versus turkey against chicken genome) could be noted and are probably caused by different qualities of the respective reference genomes. Notwithstanding, these calculated values can now be applied by the AFS user to objectively assign quantification values as potential false-positives, as done above in the case of the doner kebap samples.

#### Matrix effects and their possible correction by linear regression

Different types of food matrices can bias quantification analyses because different tissues often contain varying concentrations of DNA, and cellular DNA may also be extracted from them at different efficiencies. To study this effect, we included three types of sausage matrices: the Kal samples, consisting only of pure meat, the KLyo samples, in which pork material was represented by three tissues (meat, rind and lard at a ratio of 1:4:15) and the KGeflLyo sausages, containing chicken material as a 1:1 mixture of meat and skin (Online Tab. S1).

Specifically for the KGeflLyo sausages with their partial replacement of chicken meat by skin (Online Fig. S2), the chicken component showed a substantial overrepresentation on the DNA level, thus severely compromising the quantification results for this matrix type (independent of whether AFS or PCR methods were applied; comp. Tab. 1). While samples containing meat-only chicken showed minimal deviations of 0.1-0.5 % from expected values, the meat/skin matrix led to an almost proportional overestimation of chicken by 9-20 % (Tab. 1d; Online Fig. S2). A second, but milder effect was noticed for pork as an ingredient, which was systematically underestimated by 2.7-7.8 % in the KLyo A-D, 0.1-0.6 % in the KGeflLyo A-D and 0-5.5 % in the Kal A-E samples, respectively.

Assuming that the observed effects represent systematic errors, we decided to normalize our measurements by applying linear regression. We did this for every sample type and species separately to consider both matrix-specific and species-dependent effects (see calibrated AFS values in Tab. 1). In fact, the improvement of the quantification values turned out to be massive, showing that AFS (very much like the PCR methods; comp. [8, 13]) will benefit from the establishment of such matrix calibration factors. Indeed, we were able to correct efficiently for most of the systematic error over a broad range of expected values. Note however, that in some cases (e.g. Kal A and E) deviation slightly increased after the normalisation procedure for the very low expected values of 0.5 and 1 %, respectively (Tab. 1b).

#### Limits of detection and precision

Using normalised values gained by linear regression, we calculated the limit of detection (LoD) of AFS at a confidence level of 95%, applying the procedure described by [25]. The LoD describes the lowest quantity of an analyte that can be reliably detected above the observed background noise. In the case of read-mapping approaches, LoD will depend on genome relatedness and resulting chance for false-positive read assignment, which in turn partly depends on read length (see above). If closely related genomes (e.g. sheep vs. goat and cattle vs. buffalo) are included in an AFS mapping procedure using 150 bp reads, the method produces a LoD of 1.6 %. If only distant species are tested for, the LoD decreases to 1.0 %.

To also infer the random error produced by AFS, and thus the precision of the method, we calculated 95% confidence intervals for every instance of the expected species proportions between 0.5 and 80% (Online Tab. S3). Proportion components below 2% are measured with about 50 % uncertainty. Measurement error decreased to about 10 % for proportions between 2 and 36 % and 4 % for proportions above 36%. Overall, CIs turned out to be close to the expected values and therefore are an excellent indication of high AFS precision over the entire range of expected values from 0.5 to 80%.

## Discussion

Classical DNA-based species identification in food is routinely performed as a targeted approach using PCR-based methods, which can detect only a certain range of taxa, for which the PCR primers ideally fit [5–8, 10–13]. AFS in contrast analyses the complete DNA of a foodstuff without amplification and is therefore a non-targeted, whole-genome screening approach [18]. To investigate the potential of AFS to detect unforeseen species components, we chose to study a real-case food control scenario and sequenced the meat from five doner kebap samples from the Rhine-Main area. According to German food legislation, snacks sold under the label “doner kebap” are expected to consist only of sheep and/or beef [26]. However, occasional surveys conducted by food authorities [27] or even occasioned by broadcasting stations [28] have already pointed at a considerable heterogeneity of animal species components in doner kebap samples from Germany, which very often contained unlabelled poultry (chicken, turkey) and in rare cases even pork. Using AFS, we found that 3 of our 5 samples indeed contained turkey meat, two samples even at a major extent (90 % or more). None of the samples, however, was openly advertised to the consumer as “poultry doner”. In addition, AFS detected in four cases soy as an unexpected and unlabelled ingredient, which may be critical for consumers suffering from allergy towards soybeans. Soybean DNA may originate from the usage of spice coating (panada). One sample additionally contained maize DNA, the origin of which is unclear. AFS thus confirmed the results previously obtained by other labs in doner kebap species screens and thus should function well as a method in routine food screening.

The performance of AFS for the quantification of species in different types of food matrices has not yet been investigated systematically. The main focus of the current study was therefore to explore the quantification potential of AFS to infer species proportions of reference sausages, which have previously been used in the field to evaluate PCR-based quantification methods. To directly compare AFS to quantification results obtained by qPCR [8] and ddPCR [13], we calculated for simplicity the sum of the % deviation (measured vs. expected) for each sausage sample (Tab. 1b-d). Results showed that AFS data -very much like the qPCR and ddPCR data-need to be calibrated for matrix-dependent biases to generate the most accurate results. Indeed, in 8 of 12 cases “AFS-cal” produced the best results, while ddPCR turned out to be clearly superior in one case (Kal C) and slightly better in three cases (KLyo A, C, D). AFS readily identifies and quantifies proportions of species over a broad % range. Most importantly it works at the 1% level, a value often approximatively taken by food authorities to distinguish problematic species amounts from trace amounts, e.g. originating by unavoidable contamination.

Very much like for other DNA-based methods, the limitations of AFS are set by sequence similarities between closely related genomes and by the so-called matrix effect, which ultimately determines the extent to which species proportions in food can be indirectly inferred from DNA proportions. Our theoretical evaluation of possible wrong read assignments between closely related taxa provides the applicant of AFS with a means to readily distinguish between true and false quantification results. Food consisting of species, which have diverged at minimum 10 mio years ago (e.g. sheep-goat or cattle-buffalo), may thus be analysed without much problems. If AFS is performed for other, possibly closer taxa, the limits of false-positives can easily be determined by the procedure, which we have outlined in the methods section.

As previously noticed for PCR-based quantification methods [8, 13], the AFS requires mathematical calibration for matrix effects to achieve best results (see above). Theoretically, for instance, it should be necessary for AFS to take into account that birds have only 1/3 the genome size of mammals. In practice, this consideration proved to be not useful at all for quantifying food containing a mixture of bird and mammalian material by AFS (data not shown). The possible reason is that chicken meat may contain more DNA per gram tissue than, e.g., pork [29], thus compensating for the smaller genome size. It will be almost impossible to define the DNA amounts for all conceivable tissues from food-relevant species. However, the application of food matrix reference material, as done in the present study, facilitates a guided calibration of matrix effects and thus efficiently circumvents this problem.

In conclusion, we confirm here that AFS is a potent additional screening and quantification tool in the repertoire of foodstuff analysis. We have calculated that AFS sequencing reagent costs (50 libraries prepared in parallel, 500 k reads each, all loaded on 1 Illumina MiSeq flowcell) currently would amount to appr. 90 EUR per sample (see [19] for high-multiplex estimations). The computer skills required match those of a typical bioinformatics master student, and routine screening of 1 mio reads against up to 10 eukaryotic genomes can be performed on a laptop PC requiring a computation time of appr. 20 min (see Materials and Methods for hardware used). We like to point out that, in contrast to standard PCR analytics and depending on the desired depth of analysis, the AFS can go well beyond the mere identification of animal and plant species into the world of food microbiota, even including viruses [18]. Ideally, AFS would screen for the ever-growing number of sequenced animal, plant, fungal and bacterial genomes in one single analysis on standard computers. However, due to the usage of algorithms involving read mapping and sequence alignments, the screening power of AFS is currently limited to 20-30 species with large eukaryotic genomes in one analysis. We therefore investigate the applicability of novel non-alignment-based, memory-efficient algorithms for AFS. At the same time, the identification and quantification of microbiota from foodstuff by AFS is a goal worth of pursuing in future.

## Conflict of Interest

The authors declare that they have no conflict of interest.

## Compliance with ethics requirements

This article does not contain any studies with human or living animal subjects.

## Acknowledgements

TH and SLH gratefully acknowledge funding by the Federal Office for Agriculture and Food (project ID: 2816503814).

## References

1. German Federal Office for Risk Assessment: Food safety and globalisation - challenges and opportunities (Bundesamt für Risikobewertung). https://www.bfr.bund.de/en/press_information/2014/13/food_safety_and_globalisation__challenges_and_opportunities-190341.html. Accessed 28 Aug 2019

2. German Federal Office of Consumer Protection and Food Safety (Bundesamt für Verbraucherschutz und Lebensmittelsicherheit). https://www.bvl.bund.de/DE/Home/homepage_node.html. Accessed 28 Aug 2019

3. German Drug Law (Bundesministerium der Justiz und für Verbraucherschutz: Gesetz über den Verkehr mit Arzneimitteln). https://www.gesetze-im-internet.de/amg_1976/AMG.pdf. Accessed 28 Aug 2019

4. Swiss Food Legislation (Schweizerisches Bundesgesetz über Lebensmittel und Gebrauchsgegenstände (Lebensmittelgesetz, LMG) vom 20. Juni 2014). https://www.admin.ch/opc/de/official-compilation/2017/249.pdf. Accessed 28 Aug 2019

5. Brodmann PD, Moor D (2003) Sensitive and semi-quantitative TaqMan™ realtime polymerase chain reaction systems for the detection of beef (Bos taurus) and the detection of the family Mammalia in food and feed. Meat Sci 65:599–607. https://doi.org/10.1016/S0309-1740(02)00253-X

6. Zhang C-L, Fowler MR, Scott NW, et al (2007) A TaqMan real-time PCR system for the identification and quantification of bovine DNA in meats, milks and cheeses. Food Control 18:1149–1158. https://doi.org/10.1016/J.FOODCONT.2006.07.018

7. Köppel R, Ruf J, Zimmerli F, Breitenmoser A (2008) Multiplex real-time PCR for the detection and quantification of DNA from beef, pork, chicken and turkey. Eur Food Res Technol 227:1199–1203. https://doi.org/10.1007/s00217-008-0837-7

8. Köppel R, Ruf J, Rentsch J (2011) Multiplex real-time PCR for the detection and quantification of DNA from beef, pork, horse and sheep. Eur Food Res Technol 232:151–155. https://doi.org/10.1007/s00217-010-1371-y

9. Köppel R, Eugster A, Ruf J, Rentsch J (2012) Quantification of Meat Proportions by Measuring DNA Contents in Raw and Boiled Sausages Using Matrix-Adapted Calibrators and Multiplex Real-Time PCR. J AOAC Int 95:494–499. https://doi.org/10.5740/jaoacint.11-115

10. Ulca P, Balta H, Çağın İ, Senyuva HZ (2013) Meat species identification and Halal authentication using PCR analysis of raw and cooked traditional Turkish foods. Meat Sci 94:280–284. https://doi.org/10.1016/j.meatsci.2013.03.008

11. Floren C, Wiedemann I, Brenig B, et al (2015) Species identification and quantification in meat and meat products using droplet digital PCR (ddPCR). Food Chem 173:1054–1058. https://doi.org/10.1016/j.foodchem.2014.10.138

12. Song K-Y, Hwang HJ, Kim JH (2017) Ultra-fast DNA-based multiplex convection PCR method for meat species identification with possible on-site applications. Food Chem 229:341–346. https://doi.org/10.1016/j.foodchem.2017.02.085

13. Köppel R, Ganeshan A, Weber S, et al (2019) Duplex digital PCR for the determination of meat proportions of sausages containing meat from chicken, turkey, horse, cow, pig and sheep. Eur Food Res Technol 245:853–862. https://doi.org/10.1007/s00217-018-3220-3

14. Markoulatos P, Siafakas N, Moncany M (2002) Multiplex polymerase chain reaction: A practical approach. J Clin Lab Anal 16:47–51. https://doi.org/10.1002/jcla.2058

15. Berry D, Mahfoudh K Ben, Wagner M, Loy A (2011) Barcoded Primers Used in Multiplex Amplicon Pyrosequencing Bias Amplification. Appl Environ Microbiol 77:7846–7849. https://doi.org/10.1128/AEM.05220-11

16. Tedersoo L, Anslan S, Bahram M, et al (2015) Shotgun metagenomes and multiple primer pair-barcode combinations of amplicons reveal biases in metabarcoding analyses of fungi. MycoKeys 10:1–43. https://doi.org/10.3897/mycokeys.10.4852

17. Sze MA, Schloss PD (2019) The Impact of DNA Polymerase and Number of Rounds of Amplification in PCR on 16S rRNA Gene Sequence Data. mSphere 4:e00163–19. https://doi.org/10.1128/msphere.00163-19

18. Ripp F, Krombholz C, Liu Y, et al (2014) All-Food-Seq (AFS): a quantifiable screen for species in biological samples by deep DNA sequencing. BMC Genomics 15:639. https://doi.org/10.1186/1471-2164-15-639

19. Liu Y, Ripp F, Koeppel R, et al (2017) AFS: identification and quantification of species composition by metagenomic sequencing. Bioinformatics 33:btw822. https://doi.org/10.1093/bioinformatics/btw822

20. Eugster A, Ruf J, Rentsch J, Köppel R (2009) Quantification of beef, pork, chicken and turkey proportions in sausages: use of matrix-adapted standards and comparison of single versus multiplex PCR in an interlaboratory trial. Eur Food Res Technol 230:55–61. https://doi.org/10.1007/s00217-009-1138-5

21. FASTQC. A quality control tool for high throughput sequence data. https://www.bioinformatics.babraham.ac.uk/projects/fastqc/. Accessed 28 Aug 2019

22. Bolger AM, Lohse M, Usadel B (2014) Trimmomatic: a flexible trimmer for Illumina sequence data. Bioinformatics 30:2114–2120. https://doi.org/10.1093/bioinformatics/btu170

23. BBTools. A suite of fast, multithreaded bioinformatics tools designed for analysis of DNA and RNA sequence data. https://sourceforge.net/projects/bbmap/. Accessed 28 Aug 2019

24. Kumar S, Stecher G, Suleski M, Hedges SB (2017) TimeTree: A Resource for Timelines, Timetrees, and Divergence Times. Mol Biol Evol 34:1812–1819. https://doi.org/10.1093/molbev/msx116

25. Armbruster DA, Pry T (2008) Limit of blank, limit of detection and limit of quantitation. Clin Biochem Rev 29 Suppl 1:S49–52

26. Marking of “doner kebab” and “similar” products by bulk delivery. (Kenntlichmachung von „Döner Kebab“ und „ähnlichen“ Erzeugnissen bei loser Abgabe. Bayerisches Landesamt für Gesundheit und Lebensmittelsicherheit). https://www.lgl.bayern.de/downloads/lebensmittel/doc/merkblatt_doener_kebab.pdf. Accessed 28 Aug 2019

27. The composition and labelling of doner kebabs. LACORS. https://www.ihsti.com/lacors/ContentDetails.aspx?id=21001. Accessed 28 Aug 2019

28. Frequently minced meat and additives in doner kebab. (Häufig Fleischbrät und Zusatzstoffe im Döner. Norddeutscher Rundfunk). https://www.ndr.de/ratgeber/verbraucher/Haeufig-Fleischbraet-und-Zusatzstoffe-im-Doener,doener164.html. Accessed 28 Aug 2019

29. Cai Y, Li X, Lv R, et al (2014) Quantitative analysis of pork and chicken products by droplet digital PCR. Biomed Res Int 2014:810209. https://doi.org/10.1155/2014/810209

