## Supplementary material for "Identification and quantification of meat product ingredients by whole-genome metagenomics (All-Food-Seq)": Online Fig. S1

**(a)**

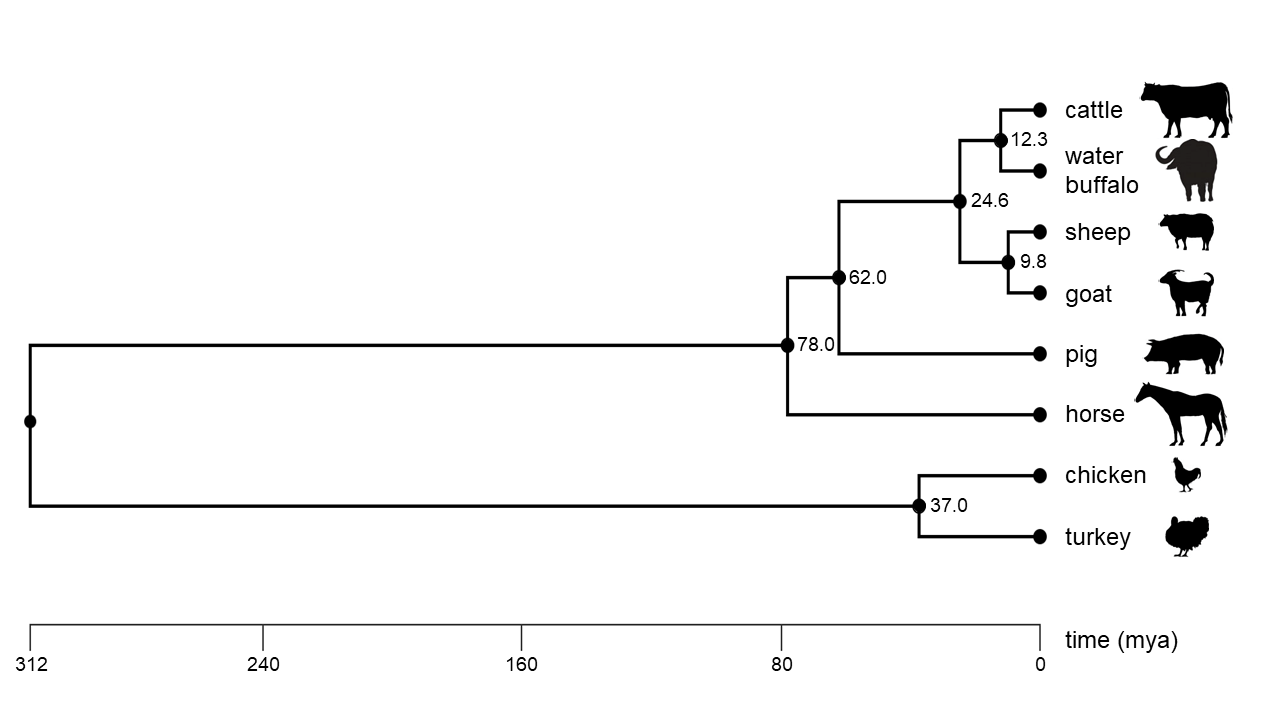

**(b)**

fp buffalo [%]

| proportion cattle [%] | 50 bp | 100 bp | 150 bp |
| --- | --- | --- | --- |
| 100 | 2.65 | 1.87 | 1.17 |
| 75 | 1.85 | 1.37 | 0.87 |
| 50 | 1.17 | 0.88 | 0.56 |
| 25 | 0.54 | 0.43 | 0.27 |
| 10 | 0.22 | 0.17 | 0.11 |
| 5 | 0.11 | 0.09 | 0.05 |
| 1 | 0.02 | 0.02 | 0.01 |

fp chicken [%]

| proportion turkey [%] | 50 bp | 100 bp |
| --- | --- | --- |
| 100 | 0.19 | 0.07 |
| 75 | 0.14 | 0.05 |
| 50 | 0.09 | 0.04 |
| 25 | 0.05 | 0.02 |
| 10 | 0.02 | 0.01 |
| 5 | 0.01 | 0 |
| 1 | 0 | 0 |

fp turkey [%]

| proportion chicken [%] | 50 bp | 100 bp | 150 bp |
| --- | --- | --- | --- |
| 100 | 0.38 | 0.09 | 0.04 |
| 75 | 0.29 | 0.07 | 0.03 |
| 50 | 0.2 | 0.05 | 0.02 |
| 25 | 0.1 | 0.02 | 0.01 |
| 10 | 0.05 | 0.02 | 0 |
| 5 | 0.03 | 0.01 | 0 |
| 1 | 0.01 | 0.01 | 0 |

fp sheep [%]

| proportion goat [%] | 50 bp | 100 bp | 150 bp |
| --- | --- | --- | --- |
| 100 | 3.05 | 2.35 | 1.69 |
| 75 | 2.06 | 1.71 | 1.26 |
| 50 | 1.24 | 1.1 | 0.83 |
| 25 | 0.56 | 0.53 | 0.41 |
| 10 | 0.21 | 0.21 | 0.16 |
| 5 | 0.11 | 0.1 | 0.08 |
| 1 | 0.02 | 0.02 | 0.01 |

fp goat [%]

| proportion sheep [%] | 50 bp | 100 bp | 150 bp |
| --- | --- | --- | --- |
| 100 | 5.08 | 3.65 | 2.68 |
| 75 | 3.41 | 2.61 | 1.96 |
| 50 | 2.03 | 1.66 | 1.26 |
| 25 | 0.92 | 0.78 | 0.61 |
| 10 | 0.35 | 0.32 | 0.25 |
| 5 | 0.18 | 0.17 | 0.13 |
| 1 | 0.04 | 0.03 | 0.02 |

**Online Fig. S1 Species phylogeny and evaluation of false-positive (fp) read assignments. (a)** Phylogenetic relationships of animal species used in this study. Data are based on http://www.timetree.org, accessed on 12 July 2019. **(b)** False- positive read assignments for closely related species pairs during AFS analysis were evaluated using AFS read mapping for sequence read lengths of 50, 100 and 150 bp. No 150 bp sequence reads were available in the NCBI SRA database for turkey and therefore could not be analysed. The x-axis gives the true proportion of species reads in a dataset the y-axis depicts the % of falsely assigned reads to the related species. Graphical representations are based on the values shown in Online Tab. S3.
