## Supplementary material for "Identification and quantification of meat product ingredients by whole-genome metagenomics (All-Food-Seq)": Online Fig. S2

beef KLyo

measured [%]

| expected [%] | expected |
| --- | --- |
| 14 | 14 |
| 36 | 36 |
| 58 | 58 |
| 80 | 80 |
| 0.5 | 0.5 |
| 2 | 2 |
| 4 | 4 |
| 5.5 | 5.5 |
| 1 | 1 |
| 9 | 9 |
| 25 | 25 |
| 35 | 35 |
| 55 | 55 |
| 100 | 100 |

| expected [%] | data |
| --- | --- |
| 14 | 19.33 |
| 36 | 41.78 |
| 58 | 66.03 |
| 80 | 82.81 |

beef KGeflLyo

measured [%]

| expected [%] | expected |
| --- | --- |
| 14 | 14 |
| 36 | 36 |
| 58 | 58 |
| 80 | 80 |
| 0.5 | 0.5 |
| 2 | 2 |
| 4 | 4 |
| 5.5 | 5.5 |
| 1 | 1 |
| 9 | 9 |
| 25 | 25 |
| 35 | 35 |
| 55 | 55 |
| 100 | 100 |

| expected [%] | data |
| --- | --- |
| 0.5 | 0.64 |
| 2 | 1.96 |
| 4 | 3.36 |
| 5.5 | 3.88 |

beef Kal

measured [%]

| expected [%] | expected |
| --- | --- |
| 14 | 14 |
| 36 | 36 |
| 58 | 58 |
| 80 | 80 |
| 0.5 | 0.5 |
| 2 | 2 |
| 4 | 4 |
| 5.5 | 5.5 |
| 1 | 1 |
| 9 | 9 |
| 25 | 25 |
| 35 | 35 |
| 55 | 55 |
| 100 | 100 |

| expected [%] | data |
| --- | --- |
| 1 | 1.35 |
| 9 | 11.19 |
| 25 | 25.83 |
| 35 | 38.23 |
| 55 | 56.88 |

pork KLyo

measured [%]

| expected [%] | expected |
| --- | --- |
| 14 | 14 |
| 36 | 36 |
| 58 | 58 |
| 80 | 80 |
| 0.5 | 0.5 |
| 2 | 2 |
| 4 | 4 |
| 5.5 | 5.5 |
| 1 | 1 |
| 9 | 9 |
| 25 | 25 |
| 35 | 35 |
| 55 | 55 |
| 100 | 100 |

| expected [%] | data |
| --- | --- |
| 80 | 74.63 |
| 58 | 52.39 |
| 36 | 28.25 |
| 14 | 11.26 |

pork KGeflLyo

measured [%]

| expected [%] | expected |
| --- | --- |
| 14 | 14 |
| 36 | 36 |
| 58 | 58 |
| 80 | 80 |
| 0.5 | 0.5 |
| 2 | 2 |
| 4 | 4 |
| 5.5 | 5.5 |
| 1 | 1 |
| 9 | 9 |
| 25 | 25 |
| 35 | 35 |
| 55 | 55 |
| 100 | 100 |

| expected [%] | data |
| --- | --- |
| 5.5 | 5.07 |
| 4 | 3.47 |
| 2 | 1.42 |
| 0.5 | 0.37 |

pork Kal

measured [%]

| expected [%] | expected |
| --- | --- |
| 14 | 14 |
| 36 | 36 |
| 58 | 58 |
| 80 | 80 |
| 0.5 | 0.5 |
| 2 | 2 |
| 4 | 4 |
| 5.5 | 5.5 |
| 1 | 1 |
| 9 | 9 |
| 25 | 25 |
| 35 | 35 |
| 55 | 55 |
| 100 | 100 |

| expected [%] | data |
| --- | --- |
| 35 | 30.9 |
| 55 | 49.52 |
| 25 | 19.75 |
| 9 | 7.71 |
| 1 | 1.02 |

chicken KLyo

measured [%]

| expected [%] | expected |
| --- | --- |
| 14 | 14 |
| 36 | 36 |
| 58 | 58 |
| 80 | 80 |
| 0.5 | 0.5 |
| 2 | 2 |
| 4 | 4 |
| 5.5 | 5.5 |
| 1 | 1 |
| 9 | 9 |
| 25 | 25 |
| 35 | 35 |
| 55 | 55 |
| 100 | 100 |

| expected [%] | data |
| --- | --- |
| 0.5 | 0.65 |
| 2 | 2.19 |
| 4 | 3.91 |
| 5.5 | 4.98 |

chicken KGeflLyo

measured [%]

| expected [%] | expected |
| --- | --- |
| 14 | 14 |
| 36 | 36 |
| 58 | 58 |
| 80 | 80 |
| 0.5 | 0.5 |
| 2 | 2 |
| 4 | 4 |
| 5.5 | 5.5 |
| 1 | 1 |
| 9 | 9 |
| 25 | 25 |
| 35 | 35 |
| 55 | 55 |
| 100 | 100 |

| expected [%] | data |
| --- | --- |
| 14 | 29.79 |
| 36 | 56.9 |
| 58 | 75.69 |
| 80 | 89.19 |

sheep Kal

measured [%]

| expected [%] | expected |
| --- | --- |
| 14 | 14 |
| 36 | 36 |
| 58 | 58 |
| 80 | 80 |
| 0.5 | 0.5 |
| 2 | 2 |
| 4 | 4 |
| 5.5 | 5.5 |
| 1 | 1 |
| 9 | 9 |
| 25 | 25 |
| 35 | 35 |
| 55 | 55 |
| 100 | 100 |

| expected [%] | data |
| --- | --- |
| 9 | 10.12 |
| 1 | 1.31 |
| 25 | 28.19 |
| 55 | 50.76 |
| 35 | 32.18 |

turkey KLyo

measured [%]

| expected [%] | expected |
| --- | --- |
| 14 | 14 |
| 36 | 36 |
| 58 | 58 |
| 80 | 80 |
| 0.5 | 0.5 |
| 2 | 2 |
| 4 | 4 |
| 5.5 | 5.5 |
| 1 | 1 |
| 9 | 9 |
| 25 | 25 |
| 35 | 35 |
| 55 | 55 |
| 100 | 100 |

| expected [%] | data |
| --- | --- |
| 5.5 | 5.25 |
| 4 | 3.35 |
| 2 | 1.35 |
| 0.5 | 0.37 |

turkey KGeflLyo

measured [%]

| expected [%] | expected |
| --- | --- |
| 14 | 14 |
| 36 | 36 |
| 58 | 58 |
| 80 | 80 |
| 0.5 | 0.5 |
| 2 | 2 |
| 4 | 4 |
| 5.5 | 5.5 |
| 1 | 1 |
| 9 | 9 |
| 25 | 25 |
| 35 | 35 |
| 55 | 55 |
| 100 | 100 |

| expected [%] | data |
| --- | --- |
| 80 | 64.49 |
| 58 | 37.64 |
| 36 | 19.51 |
| 14 | 6.54 |

horse Kal

measured [%]

| expected [%] | expected |
| --- | --- |
| 14 | 14 |
| 36 | 36 |
| 58 | 58 |
| 80 | 80 |
| 0.5 | 0.5 |
| 2 | 2 |
| 4 | 4 |
| 5.5 | 5.5 |
| 1 | 1 |
| 9 | 9 |
| 25 | 25 |
| 35 | 35 |
| 55 | 55 |
| 100 | 100 |

| expected [%] | data |
| --- | --- |
| 55 | 57.27 |
| 35 | 37.83 |
| 25 | 25.09 |
| 1 | 1.3 |
| 9 | 8.48 |

**Online Fig. S2 Evaluation of the matrix effect observed for each species and the three different sausage types.** Expected proportions and amounts measured by AFS analysis are plotted for each species and sausage type to evaluate the extent of the matrix effect. The identity line is given by the thick dashed line. The regression line derived from the data is thin and dashed. Equations for linear regression were used for calibration of AFS results to calibrate for the matrix effect. Note the differences between observed and expected values for chicken and turkey in KGeflLyo and for beef and pork in KLyo samples.
