## Supplementary material for "Identification and quantification of meat product ingredients by whole-genome metagenomics (All-Food-Seq)": Online Tab. S1

**Online Tab. S1 Calibration sausages and doner kebab samples analysed by AFS.** Meat compositions and tissue matrix is shown for the calibration material (Eugster et al. 2008; Köppel et al. 2011). “pe” and “se” designate paired-end and single-end read mode during sequencing, respectively.

| Sample | % beef | % pork | % chicken | % turkey | % horse | % sheep | tissue matrix | reference | read length (bp) | # reads |
| --- | --- | --- | --- | --- | --- | --- | --- | --- | --- | --- |
| Kal A | 1 | 35 | 0 | 0 | 55 | 9 | all meat | Köppel et al. 2011 | 150 pe | 2 x 1.697.010 |
| Kal B | 9 | 55 | 0 | 0 | 35 | 1 | all meat | Köppel et al. 2011 | 150 pe | 2 x 197.538 |
| Kal C | 25 | 25 | 0 | 0 | 25 | 25 | all meat | Köppel et al. 2011 | 150 pe | 2 x 820.260 |
| Kal D | 35 | 9 | 0 | 0 | 1 | 55 | all meat | Köppel et al. 2011 | 150 pe | 2 x 819.232 |
| Kal E | 55 | 1 | 0 | 0 | 9 | 35 | all meat | Köppel et al. 2011 | 150 pe | 2 x 587.426 |
| KLyo A | 14 | 80 | 0.5 | 5.5 | 0 | 0 | pork rind:meat:lard 1:4:15 | Eugster et al. 2008 | 150 pe | 2 x 819.914 |
| KLyo B | 36 | 58 | 2 | 4 | 0 | 0 | pork rind:meat:lard 1:4:15 | Eugster et al. 2008 | 150 pe | 2 x 614.942 |
| KLyo C | 58 | 36 | 4 | 2 | 0 | 0 | pork rind:meat:lard 1:4:15 | Eugster et al. 2008 | 150 pe | 2 x 1.033.814 |
| KLyo D | 80 | 14 | 5.5 | 0.5 | 0 | 0 | pork rind:meat:lard 1:4:15 | Eugster et al. 2008 | 150 pe | 2 x 851.816 |
| KGeflLyo A | 0.5 | 5.5 | 14 | 80 | 0 | 0 | chicken meat:skin 1:1 | Eugster et al. 2008 | 150 pe | 2 x 690.798 |
| KGeflLyo B | 2 | 4 | 36 | 58 | 0 | 0 | chicken meat:skin 1:1 | Eugster et al. 2008 | 150 pe | 2 x 783.536 |
| KGeflLyo C | 4 | 2 | 58 | 36 | 0 | 0 | chicken meat:skin 1:1 | Eugster et al. 2008 | 150 pe | 2 x 714.412 |
| KGeflLyo D | 5.5 | 0.5 | 80 | 14 | 0 | 0 | chicken meat:skin 1:1 | Eugster et al. 2008 | 150 pe | 2 x 1.647.746 |
| Doner Kebab 1 | n.a. | n.a. | n.a. | n.a. | n.a. | n.a. | n.a. | n.a. | 50 se | 1.124.615 |
| Doner Kebab 2 | n.a. | n.a. | n.a. | n.a. | n.a. | n.a. | n.a. | n.a. | 50 se | 2.643.404 |
| Doner Kebab 3 | n.a. | n.a. | n.a. | n.a. | n.a. | n.a. | n.a. | n.a. | 50 se | 2.415.054 |
| Doner Kebab 4 | n.a. | n.a. | n.a. | n.a. | n.a. | n.a. | n.a. | n.a. | 50 se | 1.562.922 |
| Doner Kebab 5 | n.a. | n.a. | n.a. | n.a. | n.a. | n.a. | n.a. | n.a. | 50 se | 520.326 |
