## Supplementary material for "Identification and quantification of meat product ingredients by whole-genome metagenomics (All-Food-Seq)": Online Tab. S3

**Online Tab. S3 95 % confidence intervals for the detection of different species proportions.**

| **Component proportion** | **95 % confidence interval** |
| --- | --- |
| 0.5 % | 0.5 ± 0.3 % |
| 1.0 % | 1.0 ± 0.7 % |
| 2.0 % | 2.0 ± 0.2 % |
| 4.0 % | 4.0 ± 0.2 % |
| 5.5 % | 5.5 ± 0.3 % |
| 9.0 % | 9.0 ± 0.8 % |
| 14.0 % | 14.0 ± 3.3 % |
| 25.0 % | 25.0 ± 2.9 % |
| 35.0 % | 35.0 ± 0.7 % |
| 36.0 % | 36.0 ± 3.5 % |
| 55.0 % | 55.0 ± 1.1 % |
| 58.0 % | 58.0 ± 3.5 % |
| 80.0 % | 80.0 ± 3.4 % |
